# HIF1-alpha expressing cells induce a hypoxic-like response in neighbouring cancer cells

**DOI:** 10.1101/266213

**Authors:** Hannah Harrison, Henry J Pegg, Jamie Thompson, Christian Bates, Paul Shore

**Affiliations:** Faculty of Biology, Medicine and Health, University of Manchester, Michael Smith Building, Oxford Road, Manchester M13 9PT, and 44 (0), Web: http://paulshorelab.org/

**Author notes:** **Joint corresponding authors** and.

**Keywords:** breast cancer, HIF-alpha, hypoxia, tumour micro-environment, co-culture

## Abstract

Hypoxia stimulates metastasis in cancer and is linked to poor patient prognosis. In tumours, oxygen levels vary and hypoxic regions exist within a generally well-oxygenated tumour. However, whilst the heterogeneous environment is known to contribute to metastatic progression, little is known about the mechanism by which heterogeneic hypoxia contributes to cancer progression. This is largely because existing experimental models do not recapitulate the heterogeneous nature of hypoxia. The primary effector of the hypoxic response is the transcription factor Hypoxia inducible factor 1-alpha (HIF1-alpha). HIF1-alpha is stabilised in response to low oxygen levels in the cellular environment and its expression is seen in hypoxic regions throughout the tumour.

We have developed a model system in which HIF1-alpha can be induced within a sub-population of cancer cells, thus enabling us to mimic the effects of heterogeneic HIF1-alpha expression.

We show that induction of HIF1-alpha not only recapitulates elements of the hypoxic response in the induced cells but also results in significant changes in proliferation, gene expression and mammosphere formation within the HIF1-alpha negative population.

These findings suggest that the HIF1-alpha expressing cells found within hypoxic regions are likely to contribute to the subsequent progression of a tumour by modifying the behaviour of cells in the non-hypoxic regions of the local micro-environment.

## Introduction

Tumour hypoxia greatly influences breast cancer progression (1-3) and is generally linked to poor prognosis in breast cancer patients due to increased proliferation, cell survival, invasive/metastatic ability, therapy-resistance and altered cancer stem cell activity (4-9). The main mediator of signalling in these poorly oxygenated areas is Hypoxia inducible factor 1-alpha (HIF1-alpha) which is stabilised at low oxygen levels activating numerous downstream pathways (10).

Areas of hypoxia form as tumours outgrow their blood supply and exist within otherwise well-oxygenated tumours. The hypoxic landscape is very dynamic and oxygen levels fluctuate meaning hypoxic areas shift as the tumour progresses and new blood supplies are formed (11). How the hypoxic tumour micro-environment contributes to cancer progression and metastasis is not fully understood and this is in part due to a lack of experimental models in which the heterogeneous nature of hypoxia can be modelled. Currently, *in vitro*, hypoxia is modelled in hypoxic incubators set at low oxygen levels (e.g. 1%) (6) or by treating cells with a chemical mimetic, such as Dimethyloxalylglycine (DMOG) or cobalt chloride (CoCl_2_) (12). These methods expose the whole population of cells to hypoxia, which is a situation that is unlikely to exist in tumours.

We have therefore developed a novel model in which heterogeneous expression of HIF1-alpha can be induced in a sub-population of cells. Uniquely, this model enables analysis of changes in HIF1-alpha expressing cells and their effect on surrounding cells in which HIF1-alpha is not expressed. Using this model we demonstrate that, in addition to changes in proliferation, gene expression and mammosphere forming cell activity in the HIF1-alpha expressing cells, significant changes also occur within the HIF1-alpha negative population. These findings demonstrate that HIF1-alpha expressing cancer cells have a profound effect on the co-cultured HIF1-alpha negative cells. Similar effects are likely to be seen in tumours with hypoxic, HIF1-alpha positive cells having influence upon the normoxic cells surrounding them. These changes to the local micro-environment may be important in disease progression and metastasis.

## Results

### Inducible HIF1-alpha cancer cell lines

To determine the effects of HIF1-alpha expressing cells on the surrounding HIF1-alpha negative cells we developed a model culture system in which HIF1-alpha can be induced in a sub-population of cells. To achieve this we fused a YFP-tagged destabilising domain (YFP-DD) to HIF1-alpha, thus enabling the stabilising effect of hypoxia on HIF1-alpha to be mimicked by the addition of an inducer molecule (13, 14). The DD fusion protein is stable in the presence of Trimethoprim (TMP) but when TMP is removed the protein rapidly degrades (Figure 1A). In this model YFP-DD was fused to a mutant form of HIF1-alpha (mHIF) which is stable in normoxia (15). This fusion was stably expressed in MCF7 (ER+) and MDA-MB-231 (ER-) breast cancer cell lines [MCF7-mHIF-YFP-DD and 231-mHIF-YFP-DD] and inducible expression was verified by Western blot for HIF1-alpha performed on nuclear extracts following 48 hours of culture with 10µM TMP (Figure 1B, Supplemental Figure 1 shows an unedited blot). Control lines were produced expressing YFP-DD only (MCF7-YFP-DD and 231-YFP-DD, data not shown).

**Figure 1.**
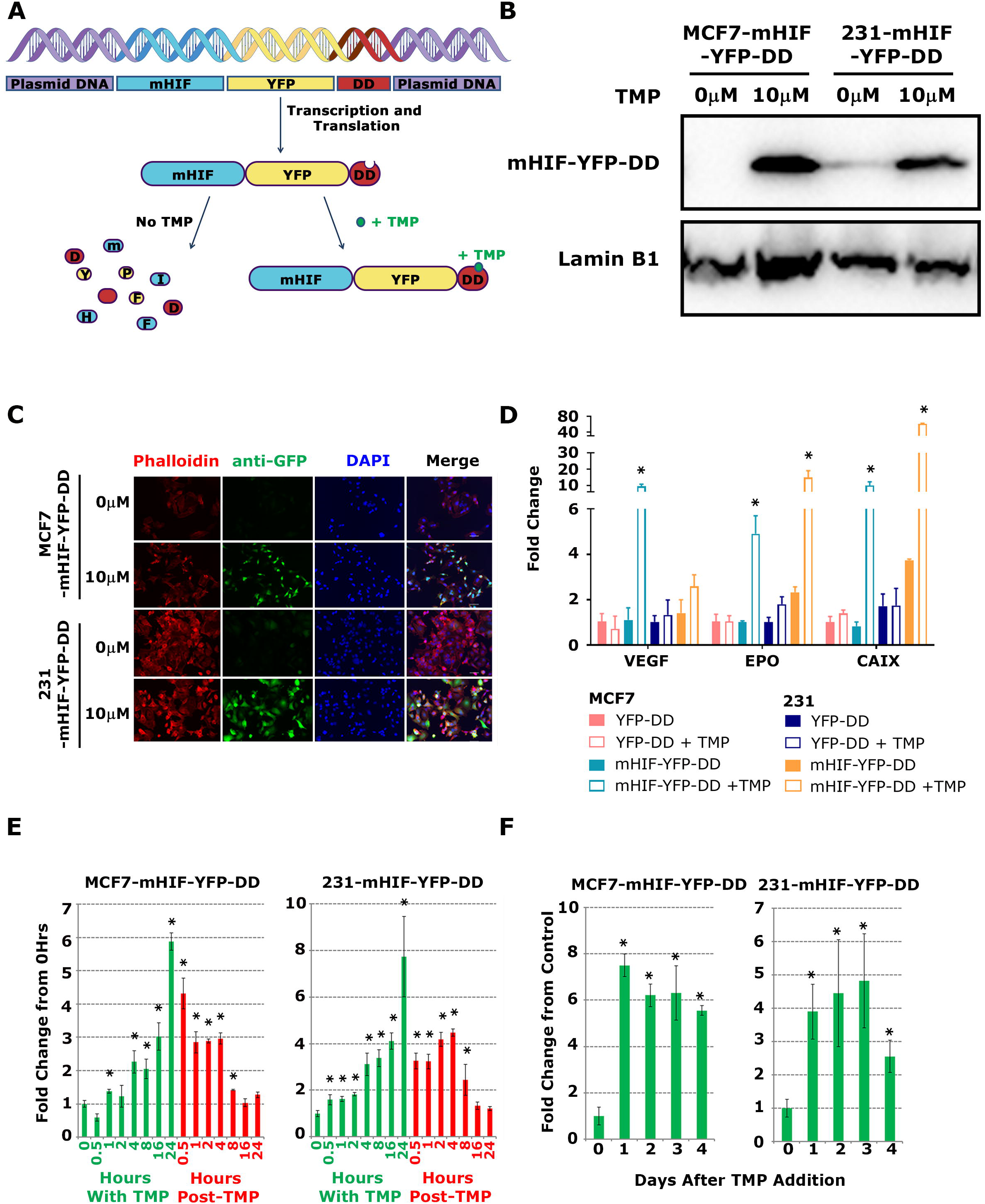
(A) Schematic detailing the inducible system used in our novel cell line model. A mutant form of HIF1-alpha (mHIF) which is stable in normoxia is fused to a YFP-tagged destabilising domain. The fusion is readily transcribed and translated but is unstable and broken down in the absence of Trimethoprim (TMP). When TMP is added the fusion is stabilised and expressed. (B) Stabilisation and expression following addition of 10µM TMP was assessed by Western blot of nuclear extracts. Western blots were performed on single membranes cut in two with upper section labelled for mHIF and lower section labelled for Lamin B1. (C) To activate downstream signalling HIF1-alpha must interact with HIF1-beta to translocate into the nucleus. Successful nuclear translocation was verified by immunocytochemistry. (D) qRT-PCR was performed on control (YFP-DD) and inducible (mHIF-YFP-DD) cells in the presence or absence of 10µM TMP. Gene expression was seen to change in the inducible cells suggesting HIF1-alpha activation of signalling. (E) Expression of mHIF-YFP-DD was measured over time following addition of TMP (green bars) and following wash out of TMP (red bars) in both MCF7 and 231 lines. Significantly increased expression was seen in both lines within 1 hour and the signal had returned to normal after 8 hours. (F) Expression of mHIF-YFP-DD was measured over time following addition of a single dose of TMP at Day 0. Expression remained elevated over 4 days.

To assess localisation and control of induction, MCF7-mHIF-YFP-DD and 231-mHIF-YFP-DD cells were cultured with 10µM TMP for 48 hours. Cells were then fixed and stained with an anti-GFP antibody to identify HIF1-alpha positive cells and Phalloidin-TRITC and DAPI to define the cytoplasm and nucleus respectively. Nuclear localisation, which is required for active HIF1-alpha signalling, was not affected by the increased size resulting from fusion to YFP-DD as the protein was present in the nucleus of induced cells (Figure 1C). Activation of HIF1-alpha target genes was subsequently determined using qRT-PCR for three known genes. Significant increases in VEGF, EPO and CAIX were seen following the addition of 10µM TMP (Figure 1D). No change was observed in the control lines. Induction occurred quickly with a significant increase in expression after 1 hour in MCF7 cells and 30 minutes in MDA-MB-231. Expression reached its highest level by 24 hours in both lines (Figure 1E). Following removal and wash out of TMP, expression rapidly decreased returning to un-induced levels after 8 hours. If TMP was not removed, induction remained high over 4 days following a single treatment with 10µM TMP (Figure 1F). Taken together these data show that the stabilising effect of hypoxia on HIF1-alpha can be mimicked by the addition of TMP in these cell lines.

### Stabilising HIF1-alpha with TMP mimics known hypoxic responses

We next determined if stabilising HIF1-alpha by the addition of TMP simulated the response known to be induced by hypoxia (1% O_2_) and a chemical hypoxia mimetic (100µM CoCl_2_). qRT-PCR was performed on a panel of known HIF1-alpha target genes (VEGF, GLUT1, EPO, CAIX and TGFA) after 48 hours induction with TMP, hypoxia or CoCl_2_. In all three conditions target genes were induced to similar levels (Figure 2A). TMP had no effect on gene expression (data not shown).

**Figure 2.**
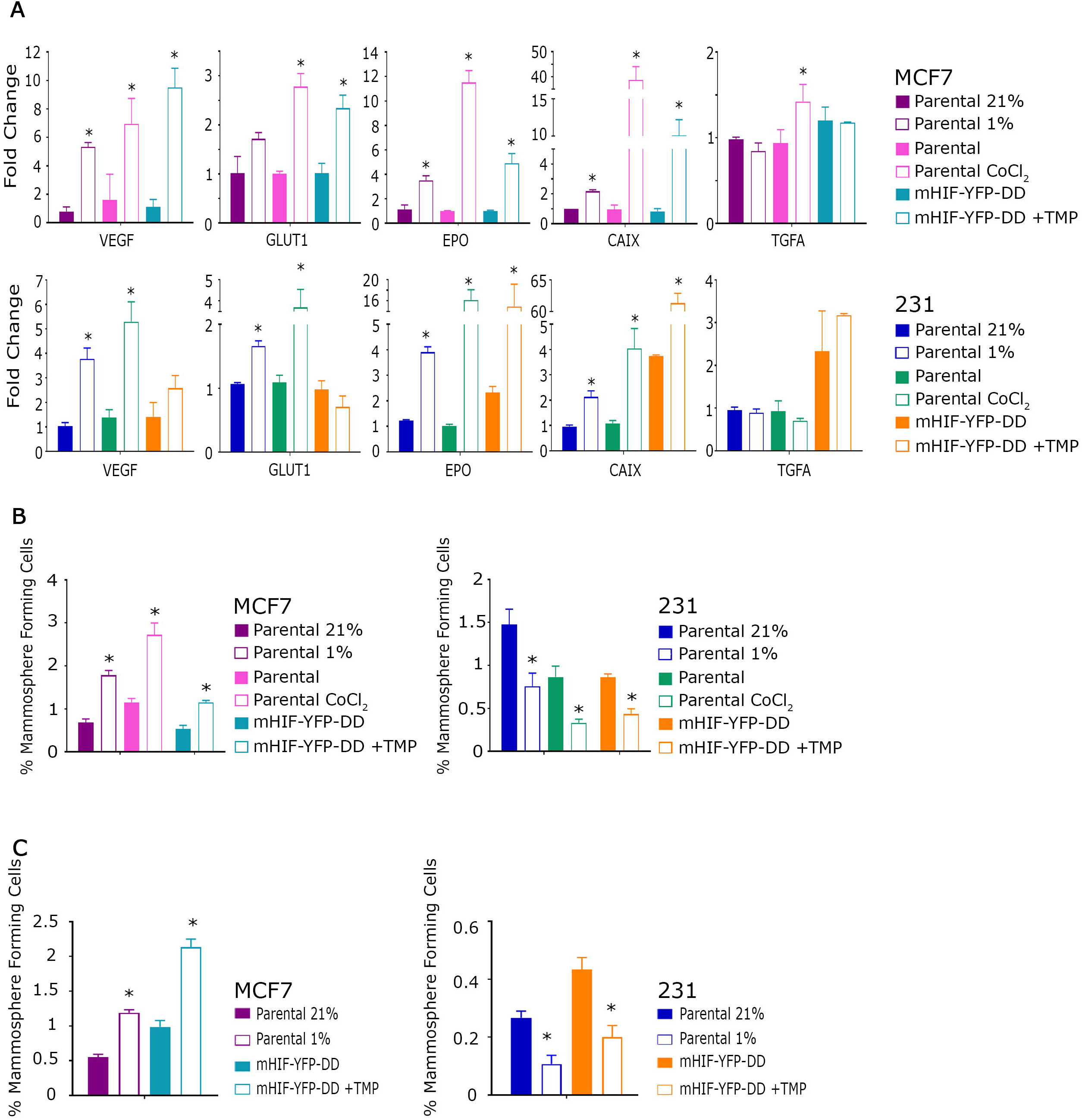
(A) qRT-PCR was performed for cells from hypoxic culture (parental 21% vs 1% O_2_), hypoxia mimetic culture (parental +/-CoCl_2_) and our HIF1-alpha inducible system (mHIF-YFP-DD +/-TMP). A similar pattern was seen in all conditions. (B) Following 48 hours in hypoxic culture (parental 21% vs 1% O_2_), hypoxia mimetic culture (parental +/-CoCl_2_) or our HIF1-alpha inducible system (mHIF-YFP-DD +/-TMP) cells were plated in mammosphere culture. A comparable response was seen in all conditions. (C) Conditioned medium was collected from cells cultured in hypoxia (parental 21% vs 1% O_2_) or our HIF1-alpha inducible system (mHIF-YFP-DD +/-TMP) and was used to treat parental cells for 48 hours. These cells were then plated in mammosphere culture. The same effect is seen in each condition.

Hypoxic culture is known to have a contrasting effect on the cancer stem cell (CSC) population in breast cancer depending upon the disease sub-type (6). In breast cancers which express estrogen receptor (ER), CSC activity increases following exposure to hypoxia. In contrast, there is a significant decrease in CSC activity in ER-negative tumours. This effect was previously shown to be directly down-stream of HIF1-alpha as siRNA targeting HIF1-alpha blocked the effect of hypoxic culture completely (6). To assess whether the same effects were seen in our HIF1-alpha expressing cells, mammosphere formation, which can be used as a measure of CSC number, was assessed following 48 hours of culture with TMP. Mammosphere number was increased in MCF7-mHIF-YFP-DD cells and decreased in 231-mHIF-YFP-DD cells as is seen in hypoxic culture and following CoCl_2_ exposure (Figure 2B).

Secreted factors are known to play a major role in the regulation of hypoxia in tumour signalling both locally and to distant sites (16). We aimed, therefore, to assess whether the CSC response seen following hypoxic culture or following HIF1-alpha induction in our model was controlled via secreted signals. Conditioned medium (CM) taken from cells cultured in hypoxia had the same effect on mammosphere formation as hypoxic culture itself; CM taken from MCF7 (ER+) cells caused an increase in mammosphere number in MCF7 whilst CM taken from MDA-MB-231 (ER-) caused a decrease in mammosphere formation in MDA-MB-231 cells (Figure 2C). This effect was blocked with addition of the HIF1-alpha inhibitor YC-1 during production of CM (Supplemental Figure 2). The same paracrine effects were seen when using CM collected from inducible lines; CM collected from MCF7-mHIF-YFP-DD and 231-mHIF-YFP-DD treated with TMP resulted in altered mammosphere formation in MCF7 and MDA-MB-231 (Figure 2C). This suggests a paracrine mediation of the signal and the protein(s) responsible for the response are secreted from the HIF1-alpha expressing cells.

### Expression of HIF1-alpha in a sub-population of cancer cells induces significant changes in surrounding cells

To simulate the heterogeneous nature of HIF1-alpha expression that is seen within a population of tumour cells we generated co-cultures by mixing wild-type MCF7 or MDA-MB-231 cells with HIF1-alpha inducible cells. After 48 hours cells were isolated by flow sorting to separate HIF1-alpha positive (YFP positive) from negative cells (Supplemental Figure 3). These cultures were subsequently used to examine the effects of the HIF1-alpha positive cells on the surrounding parental sub-population.

**Figure 3.**
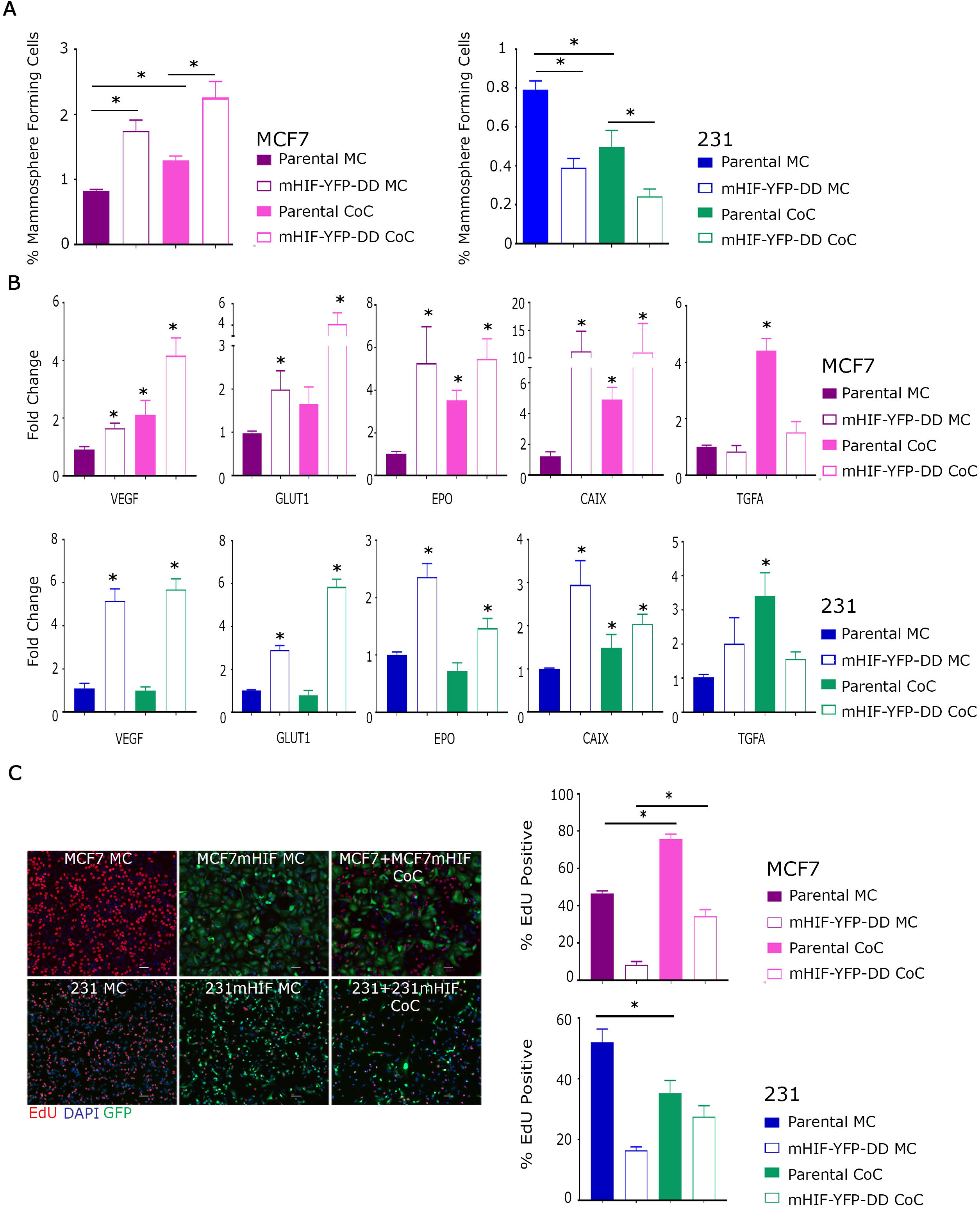
Following 48 hours in the presence of TMP in either mono-culture (MC) or co-culture (CoC), mHIF-YFP-DD and parental cells were plated into mammosphere culture or used to make RNA for qRT-PCR. (A) Mammosphere formation increased or decreased in the mHIF-YFP-DD expressing lines as expected in MCF7 and 231 cells respectively. Interestingly, mammosphere forming cell number was also affected in the both parental cell lines following co-culture. (B) Gene expression changes were seen within the HIF1-alpha inducible cells as expected. Changes were also visible, however, in the parental cells grown in co-culture. (C) Representative photomicrographs showing EdU stain in mono- and co-culture, Green anti-GFP, Blue DAPI and Red EdU. Proliferation, assessed by counting cells positive for EdU, was increased in mHIF-YFP-DD and parental cells cultured in co-culture.

Following co-culture and separation, MCF7 and MDA-MB-231 cells were plated into non-adherent culture to assess the effects on mammosphere formation. Induction of HIF1-alpha in mono-culture resulted in increased and decreased mammosphere formation as expected in MCF7 and MDA-MB-231 respectively (Figure 3A). Importantly we observed that TMP induction of HIF1-alpha in MCF7 co-cultures also resulted in a significant increase in mammosphere formation in the non-induced sub-population of cells (Figure 3A). In MDA-MB-231 co-cultures a reduction in mammospheres was observed. Moreover, qRT-PCR analysis of the HIF1-alpha target genes also revealed a significant increase in their expression in the parental, co-cultured cells (Figure 3B). This included an increase in TGF-alpha expression in the parental cell lines following co-culture, a change which is not seen in the HIF1-alpha positive inducible cells or in those cultured in hypoxia (1%) or CoCl_2_ culture. All genes responded similarly to HIF1-alpha induction in the HIF1-alpha expressing cells in both mono-and co-cultures (Figure 3B). To determine the effects of HIF1-alpha induction on the proliferative capacity of neighbouring cells, ClickIT Edu was used to assess changes in proliferation after HIF1-alpha induction in co-cultured cells. HIF1-alpha negative cell proliferation was altered in MCF7 and MDA-MB-231 when grown in co-culture with HIF1-alpha expressing cells; MCF7 cells proliferated more quickly whilst MDA-MB-231 cells grew more slowly (Figure 3C). In both lines the HIF1-alpha expressing cells proliferated more quickly when grown in co-culture compared to mono-culture suggesting a bi-directional effect of proliferation.

Taken together, these data reveal that induction of HIF1-alpha expression in a sub-population of cancer cells has a significant effect on gene expression, the mammosphere forming population and the proliferative capacity of the surrounding cancer cells. Thus our model system has revealed a complex interplay between HIF1-alpha positive and negative cancer cells that are likely to have a significant effect on the development of a tumour.

## Discussion

The inducible system described here allows HIF1-alpha, the main regulator of hypoxia, to be switched on and off within a sub-population of cells. This system has enabled us to gain a more realistic view of its effects on cancer cells within a tumour.

Using a co-culture model we have demonstrated that expression of HIF1-alpha is sufficient to induce the same expression changes in important target genes as commonly observed in hypoxia and CoCl_2_ exposure. Also, the induction of HIF1-alpha causes the same contrasting effect on mammosphere forming cells in MCF7 and MDA-MB-231 cells that result from hypoxic and CoCl_2_ culture (6). This model system revealed that when cultured together with parental cells, there is a significant effect on mammosphere formation in this HIF1-alpha negative population. Cancer stem cell (CSC) activity, measured using mammosphere culture (17), increases in MCF7 cells and decreases in MDA-MB-231 cells when cultured in the vicinity of HIF1-alpha expressing cells. This is potentially of great importance as it suggests that the CSC population in tumours may be altered both within and outside of the hypoxic zones meaning the effects of hypoxia could be far more wide reaching than initially assumed. These HIF1-alpha induced changes in the parental cell population could not have been identified using hypoxic chambers and chemical mimetics.

Our model allows us to separate and investigate the effects of the main hypoxia regulator, HIF1-alpha, from other factors which are involved in the hypoxic response within tumours, expression of HIF2-alpha for example. This allows us to see more clearly the effects of HIF1-alpha and may allow us to further elucidate the role of this protein in the hypoxic and non-hypoxic zones of a tumour. Studying the effects of these other factors is of course important and work is underway to investigate them in our model system.

Co-culture of our inducible cells with the parental lines also showed gene expression changes within the parental HIF1-alpha-negative cells. Changes included an increase in TGF-alpha expression which was not seen in the HIF1-alpha positive inducible cells or in those cultured in hypoxia (1%) or with CoCl_2_. TGF-alpha is a known target of HIF1-alpha (18) but it was not activated within our HIF1-alpha induced or hypoxic cultured cells. TGF-alpha has been previously suggested as a marker of poor outcome at follow-up and specifically for lymph node metastasis (19), whether this is linked to hypoxia is yet to be seen but this kind of change in the bulk of the tumour may explain how HIF1-alpha induction by hypoxia in the primary tumour can increase metastasis even though this recurrence often happens many years after removal of the primary tumour.

Proliferation rates were seen to be affected in both the parental and HIF1-alpha expressing cells during co-culture experiments. This bi-directional signalling cannot be identified using conventional culture systems. This data further suggests that reduced oxygen may affect not only the cells within the hypoxic region, which are expressing HIF1-alpha, but also the bulk of the tumour and supports the need for a culture system in which HIF1-alpha negative cells are represented alongside HIF1-alpha expressing cells as this change would not be seen in conventional culture.

This new inducible co-culture system allows HIF1-alpha induction within a sub-population of cancer cells and so facilitates the analysis of the complex interplay between HIF1-alpha positive and negative cells, which are found within hypoxic and normoxic tumour cells. We also note that stabilisation of HIF1-alpha is fast and the ligand can be washed out of the system with expression falling quickly. Thus the system also allows the effects of the dynamic changes in HIF1-alpha seen in tumour hypoxia to be analysed. This is important as it will allow us to model the fluctuating hypoxia observed in tumours (11). Finally, co-culture with other cell types from the local micro-environment is also possible and this may reveal more complexity in the hypoxia induced response to HIF1-alpha in tumours.

## Conclusions

This model offers the opportunity to study, in detail, the complex interactions between HIF1-alpha expressing cells and those around them. This may offer insight into the effects HIF1-alpha expressing cells have on their local micro-environment leading to altered metastasis and disease progression. Our data shows a clear bi-directional effect within the co-culture environment which could not be identified previously.

This unique model can easily be adapted to produce inducible co-cultures with other genes, which are important in other environments and diseases, meaning it offers a great deal of potential to the wider scientific community.

## Materials and Methods

### Cell lines

MCF7 and MDA-MB-231 were purchased from American Type Culture Collection. Lines were authenticated by multiplex-PCR assay using the AmpF/STR system (Applied Biosystems) and confirmed as mycoplasma free. Monolayers were grown in complete medium (DMEM/10% FCS/2 mmol/L L-glutamine/PenStrep) and maintained in a humidified incubator at 37°C at an atmospheric pressure of 5% (v/v) CO_2_/air.

### Inducible cell line production

YFP-DD was removed from pBMN YFP-DHFR(DD) (a gift from Thomas Wandless, Addgene plasmid # 29326) and cloned into HA-HIF1-alpha P402A/P564A-pcDNA3 (a gift from William Kaelin, Addgene plasmid # 18955). The new vector, mHIF-YFP-DD-pcDNA3 was used to transfect MCF7 and MDA-MB-231 using Lipofectamine 2000 according to manufacturer’s instructions. Following selection with G418 cells were FACS sorted as single cells to produce stable, clonal cell lines. Trimethoprim (TMP, Cay-16473, Cayman) was used to induce stabilisation of the fusion.

### Hypoxic and hypoxia mimetic cell culture

Cells we incubated for 48 hours in the SCI-tiveN hypoxic workstation (Ruskinn) in 1% O_2_, 5% CO_2_ and 94% N_2_ in a humidified environment at 37°C. Cells were plated, cultured, and harvested within the workstation to maintain hypoxia at all times. For cobalt chloride (CoCl_2_) induction, 100µM CoCl_2_ was added to fresh culture medium and cells were incubated for 48 hours in a humidified incubator at 37°C at an atmospheric pressure of 5% (v/v) CO_2_/air.

### Conditioned Medium

Conditioned medium (CM) was collected and spun at 1000g for 10 minutes to remove cell debris. CM was stored at −20°C for no longer than 2 weeks. CM was mixed 50:50 for fresh complete medium before being used to treat cells. 10 mmol/L YC-1 (Cayman Chemicals) was added to monolayer culture at time of plating.

### Mammosphere culture

Mammosphere culture was carried out as previously described (17), and spheres greater than 50µm were counted on day 5.

### Co-culture

Parental and inducible cells were mixed at a ratio of 50:50 and cultured as normal for 48 hours. Cells were then separated based upon expression of YFP using the FACS Aria. When comparing to mono-culture all cells were passed through the sorter to account for any pressure/sorting effect.

### Proliferation assay

Cells were plated on glass cover slides for 24 hours. ClickIT™ EdU kit Plus EdU Alexa Fluor™ 647 Imaging Kit (C10640, Thermo) was used according to manufacturer’s instructions. Briefly cells were cultured with EdU for 45 minutes before fixing, permeablising and labelling with ClickIT-647.

### Nuclear/Cytoplasmic separation

Cells were resuspended in 400μl of ice cold Buffer A (10 mM HEPES pH 7.9, 10 mM KCl, 0.1 mM EDTA, 0.1 mM EGTA, 1 mM DTT, 0.5 mM PMSF) with the addition of complete mini-EDTA-free protease inhibitor cocktail (Roche) and incubated at 4° for 15 min. Cells were lysed with 10% NP-40 (Sigma) before centrifuging at 4° and removal of the cytoplasmic extracts in the supernatant. The pellet was then resuspended in ice cold Buffer B (20 mM HEPES pH 7.9, 0.4 M NaCl, 1 mM EDTA, 1 mM EGTA, 1 mM DTT and 1 mM PMSF) containing protease inhibitors and vortexed vigorously for 45 min at 4°C. Nuclear proteins were collected from supernatant following centrifugation at 4°C. Both the nuclear and cytoplasmic extracts were stored at −80°C for future use.

### Immunocytochemistry

Cells were grown in monolayer on coverslips for 48 hours +/-10µM Trimethoprim (TMP). Medium was removed and cells were fixed and permeablised with 4% formaldehyde and 0.1% Triton for 20 minutes at room temperature. Non-specific binding was blocked using 1% BSA before antibody staining: rabbit anti-GFP (ab290, Abcam), Phalloidin-TRITC (P1951, SIGMA) or Phalloidin-iFluor 647 (ab176759, Abcam), Goat anti-Rabbit Alexa Fluor 488 (A11008, Thermo) and Goat anti-Mouse Cyanine3 (A10521, Thermo). Coverslips were mounted with ProLong™ Gold Antifade Mountant with DAPI (P36940, Thermo).

### Image analysis

Images were collected on a Zeiss Axioimager.D2 upright microscope using a 10x objective and captured using a Coolsnap HQ2 camera (Photometrics) through Micromanager software v1.4.23. Specific band pass filter sets for DAPI, FITC and Cy5 were used to prevent bleed through from one channel to the next. Images were then processed and analysed using Fiji ImageJ (http://imagej.net/Fiji/Downloads) (20) which is freely available online.

### Western blotting

Protein was separated on an SDS–PAGE and transferred to Hybond-C Extra nitrocellulose membrane. Primary antibodies included: HIF-1a (610959, BD Biosciences) and Lamin B1 (ab16048, Abcam). Densitometry was conducted using ImageJ software, which is freely available at http://rsb.info.nih.gov/ij/.

### Quantitative Reverse Transcription PCR

RNA was extracted using the Qiagen RNAeasy kit according to manufacturer’s instructions and quantified on the Nanodrop spectrophotometer (Thermo). Real time one step qRT-PCR was carried out using the QuantiTect SYBR^®^ Green RT-PCR Kit (Qiagen) according to manufacturer’s instructions before analysis on the 7900 PCR machine (Applied Biosystems).

### Statistical analysis

Data is represented as mean +/-SEM. Statistical significance was measured using parametric testing, assuming equal variance, in the majority of experiments with standard t tests for 2 paired samples used to assess difference between test and control samples.

## Acknowledgements

The Bioimaging Facility microscopes used in this study were purchased with grants from BBSRC, Wellcome and the University of Manchester Strategic Fund. Special thanks go to Roger Meadows and Steven Marsden for their help with the microscopy.

The FACS Aria within the flow cytometry core was purchased with funding from the MRC. Thanks to Michael Jackson for guidance and assistance with sorting.

