## Supplementary Materials for "HIF1-alpha expressing cells induce a hypoxic-like response in neighbouring cancer cells"

### A novel model of heterogeneous hypoxia reveals an hypoxic-like response in normoxic cancer cells

### Harrisonetal_SupplementalFigure1

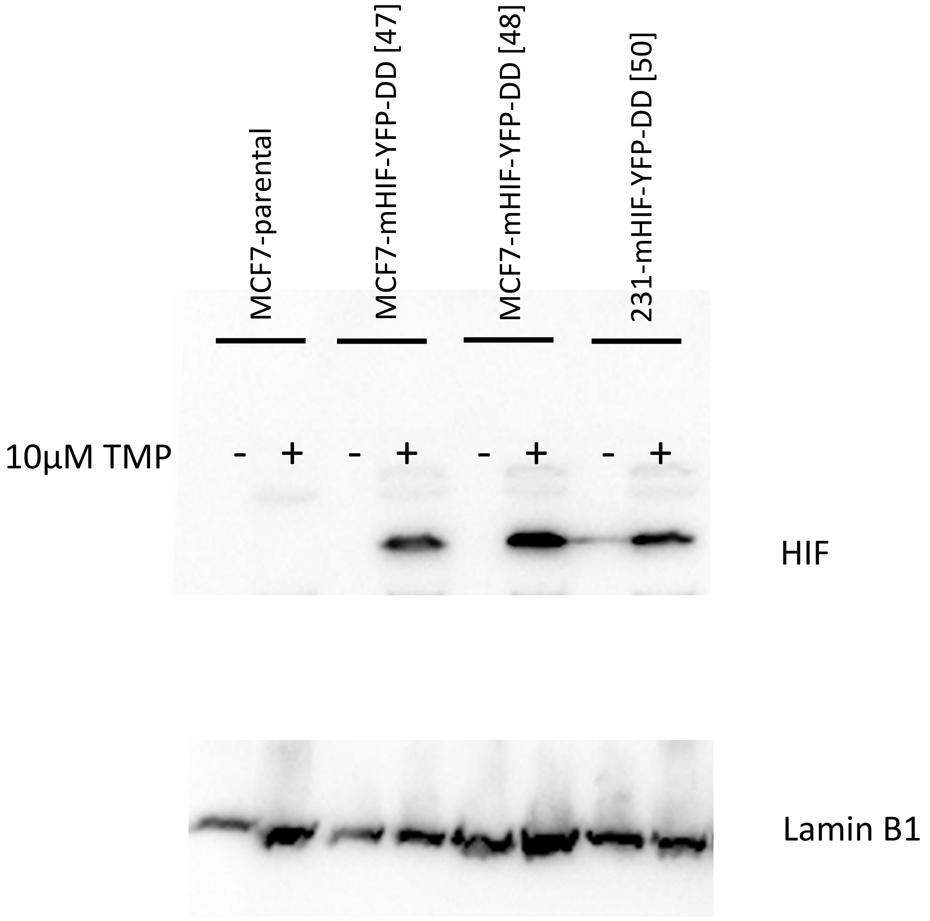


Western blot was cut in 2 halves with top being labelled for HIF and bottom for Lamin B1. This unedited western shows MCF7 parental cells, 2 MCF7-mHIF clones (#47 and 48) and 1 231-mHIF clone (#50)

### Harrisonetal_SupplementalFigure2

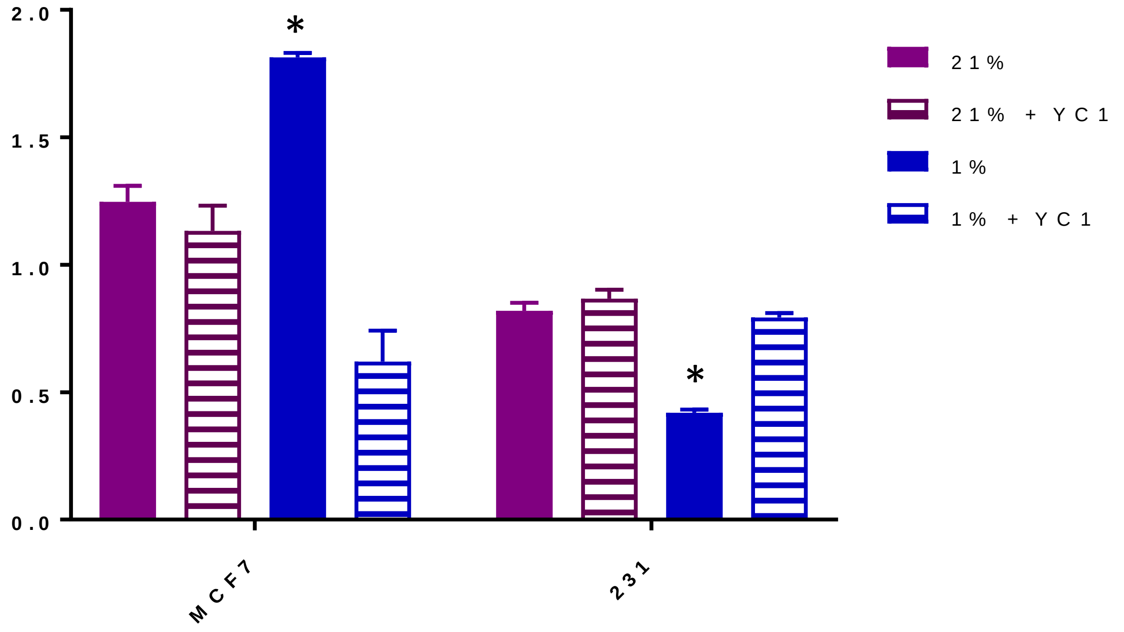


The hypoxic effect of conditioned medium taken from MCF7 and MDA-MB-231 upon mammosphere formation is blocked by HIF1 inhibitor, YC-1. Represented as mean +/- SEM. * $P<0.05$ from 21% control

### Harrisonetal_SupplementalFigure3

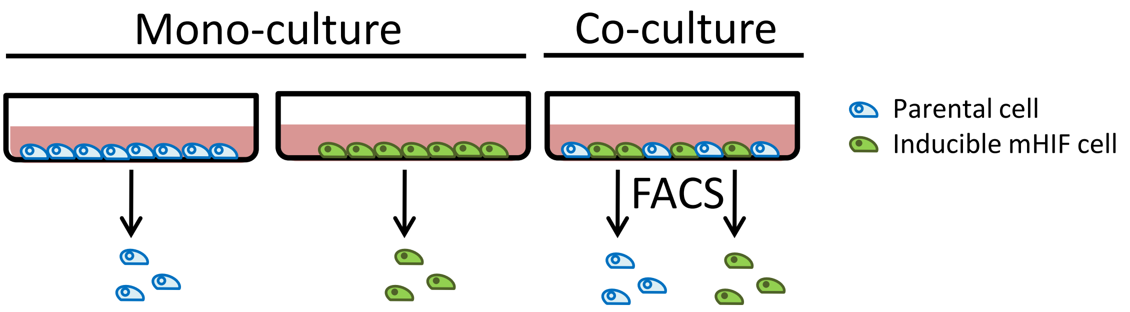


Schematic showing co-culture. Parental and inducible cells were grown either in mono-culture or as co-culture for 48 hours. The co-culture was then separated by FACS into GFP +ve and –ve populations. Mono-cultures were also exposed to FACS as a control for pressure.
